# Selective corticostriatal plasticity during acquisition of an auditory discrimination task

**DOI:** 10.1101/010066

**Authors:** Qiaojie Xiong, Petr Znamenskiy, Anthony M Zador

## Abstract

Perceptual decisions are based on the activity of sensory cortical neurons, but how organisms learn to transform this activity into appropriate actions remains unknown. Projections from the auditory cortex to the auditory striatum carry information that drives decisions in an auditory frequency discrimination task1. To assess the role of these projections in learning, we developed a Channelrhodopsin-2-based assay to selectively probe for synaptic plasticity associated with corticostriatal neurons representing different frequencies. Here we report that learning this auditory discrimination preferentially potentiates corticostriatal synapses from neurons representing either high or low frequencies, depending on reward contingencies. We observed frequency-dependent corticostriatal potentiation *in vivo* over the course of training, and *in vitro* in striatal brain slices. Our findings suggest a model in which selective potentiation of inputs representing different components of a sensory stimulus enables the learned transformation of sensory input into actions.

---

Animals use sensory information to guide their behavior. The neural mechanisms underlying the transformation of sensory responses into motor commands have been studied extensively using a two alternative forced choice (2AFC) paradigm, in which subjects are trained to make a binary decision and indicate their choice by performing one of two actions. Defined brain areas have been implicated in the circuit performing this transformation in primates^2,3^ and rodents^1,4-10^.

The site or sites of plasticity engaged when animals learn to make appropriate decisions about sensory stimuli are not well established. Striatal plasticity has been implicated in reinforcement learning^11,12^, specifically at corticostriatal inputs^13,14^. Different areas of the striatum receive input from different areas of the cortex and play different roles in learning. During motor skill learning, for example, changes occur first in the dorsomedial striatum and later in the dorsolateral striatum^14^. However, little is known about precisely what features of the cortical representation of the task are selected for potentiation in the striatum. We therefore set out to study the role of corticostriatal plasticity in task acquisition, in a setting where we could exploit our understanding of the relevant cortical representation.

We previously found that neurons in the primary auditory cortex that project to the auditory striatum drive decisions in a 2AFC auditory task^1^ in which rats learn to associate the frequency of a complex auditory cloud-of-tones stimulus with either a left or right reward port (Fig. 1a&b). Here, to test whether acquisition of this task modified the strength of corticostriatal connections, we developed an *in vivo* recording paradigm with which we could monitor corticostriatal synaptic strength in single animals over multiple behavioral sessions during the course of learning. We first injected an adeno-associated virus expressing Channelrhodopsin-2 (AAV-ChR2-Venus) into the left primary auditory cortex. This resulted in widespread expression of ChR2 in different cell types in the auditory cortex, including corticostriatal neurons and their axons in the striatum (Fig. S1). We next implanted bundles of optical fibers and tetrodes into the left auditory striatum (Fig. 1c). Brief pulses of blue light delivered through the optical fiber excited the corticostriatal axons and elicited excitatory postsynaptic responses in the striatum (Fig. 1d). Because the striatum, like the CA1 region of the hippocampus, lacks recurrent excitatory connections, we reasoned that this *in vivo* ChR2-evoked local field potential response (ChR2-LFP) could serve as a measure of the strength of the corticostriatal synaptic connectivity ^15^. The ChR2-LFP had a stereotypic waveform consisting of an early and a late component (Fig. S2). Local pharmacological blockade of excitatory but not inhibitory transmission diminished the late component (n=2 rats), indicating that it was mainly mediated by currents elicited by glutamatergic release from corticostriatal terminals (Fig. 1d, S3). The early component was resistant to all blockers including tetrodoxin, suggesting that it is driven directly by light-evoked ChR2 currents in corticostriatal axons. The early component was not observed when the electrode was advanced through the overlying somatosensory cortex, which lacked ChR2-positive axons (Fig. S4), indicating that it was not due to a photoelectric artifact. We therefore used the amplitude of this early component as a measure of the number of ChR2-expressing axons recruited by photostimulation. In subsequent analyses we normalized the ChR2-LFP to the amplitude of this presynaptic component (Fig. S2) and then used the initial slope of the second component as a measure of corticostriatal synaptic efficacy (Fig. 1d & Fig. S2b, *dotted line*). This metric was robust to changes in light intensity *in vivo* (Fig. S5) and *in vitro* (Fig. S6b) and was monotonically related to the magnitude of the intracellular EPSC when synaptic strength was manipulated experimentally *in vitro* (Fig. S6).

**Figure 1.**
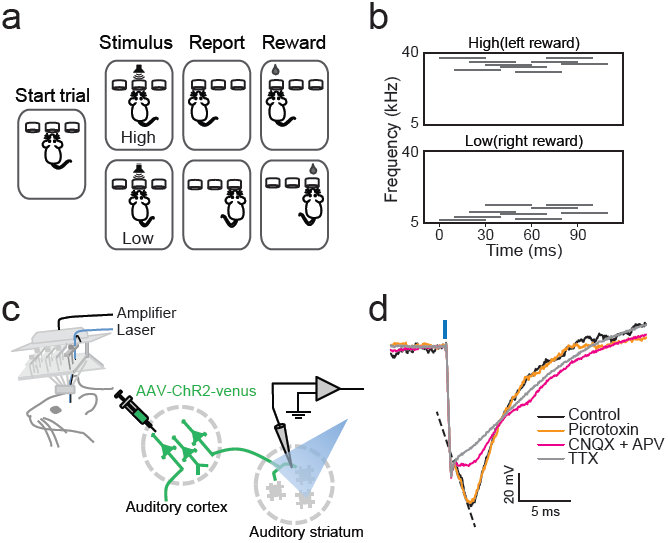
*Dissection of ChR2-LFP in vivo.* (**a**) Cloud-of-tones task. **(b)** Example spectrograms of cloud-of-tones stimuli. (**c**) Recording paradigm to examine corticostriatal synaptic strength *in vivo*. (**d**) ChR2-LFP recorded from auditory striatum under control conditions (black trace) and after application of picrotoxin (orange), CNQX and APV (pink) and tetrodotoxin (light gray). The slope of the CNQX/APV-sensitive component was used to quantify corticostriatal synaptic strength (*dotted line*).

We used the ChR2-LFP recording paradigm to assess changes in the strength of corticostriatal synapses over the course of training in the cloud-of-tones task. We first established that the ChR2-LFP signal was stable over several days in naïve rats, and then measured the ChR2-LFP after each training session. We used the tone-evoked multiunit responses recorded prior to training to estimate the frequency tuning at each site^1^ (Fig. S7, see Methods). At some recording sites, corticostriatal synaptic efficacy increased as soon as the animal started to learn the task, and continued to increase in subsequent training sessions (Fig. 2a). Synaptic efficacy at such sites thus reflected behavioral performance over the course of training. At other sites, however, corticostriatal efficacy remained unchanged over the course of training (Fig. 2b). We found that potentiation was restricted to sites tuned to low-frequency (<14 kHz, the center frequency of the tones used in the task) sounds (mean potentiation of 30%; n=17, *p*=0.002, signed-rank test, Fig. 2c), whereas sites tuned to high-frequency (>14 kHz) sounds showed no significant change (mean change of -3%; n=6, *p*=0.58, signed-rank test, Fig. 2c.). Notably, all animals in this cohort were trained to associate low frequency sounds with rightward choices (LowRight), and all recordings were performed in the left striatum. Hence, low frequencies were always associated with responses contralateral to the recording hemisphere. Our results therefore suggested that task training selectively enhances the strength of corticostriatal synapses only when the stimuli they encode are associated with contralateral choices (Fig. 2d).

**Figure 2.**
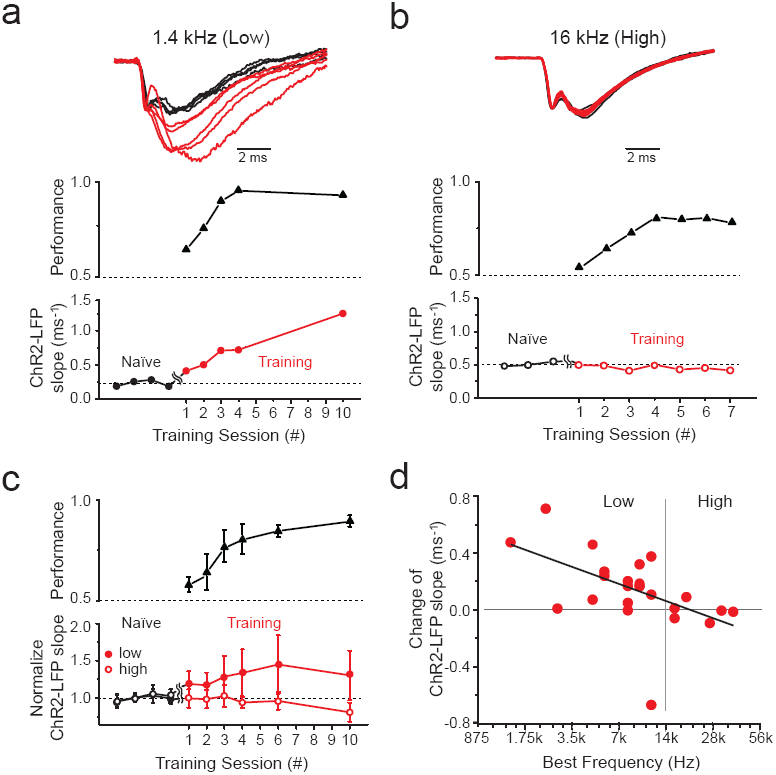
*Frequency-selective potentiation of corticostriatal ChR2-LFP slope during learning.* **(a-b)** ChR2-LFP responses (LFP slope; see Methods) before (black) and during (red) training at example sites tuned to low (**a**) and high frequency **(b)**. Session 1 is defined as the first session in which the animal performed the full task (see Methods). (**c**) Population average of normalized (see Methods) ChR2-LFP slope during learning for sites tuned to low (<14 kHz, n=16 sites, closed circles) and high (>14 kHz, n=6 sites, open circles) frequencies. Mean ± s.e. (**d**) Potentiation is restricted to sites tuned to low (<14 kHz) frequencies (23 recording sites from 8 rats; least squares regression of potentiation against frequency, *p*=0.011).

The observed potentiation depended strongly on the frequency tuning of the recording site, suggesting that corticostriatal plasticity encodes the association of specific frequencies with rewarded actions. However, since the striatum has been widely implicated in motor learning^14^, we sought to rule out this and other alternative causes of plasticity unrelated to auditory discrimination. We trained animals to perform a simple 2AFC visual task, relying on the same sequence of movements, and monitored the strength of auditory corticostriatal synapses during learning (Fig. 3a). There was no significant change in ChR2-LFP in the auditory striatum during visual task training (mean change of -17%, n=12, *p*=0.13, signed-rank test), and there was no correlation between potentiation and the preferred frequency at the recording site (n=12, *p*=0.19; Fig. 3d). However, corticostriatal inputs at these same recording sites were potentiated when the animals subsequently learned the auditory cloud-of-tones task (mean potentiation of 36%, n=6, *p*=0.03, signed-rank test; Fig. 3b&c). We therefore conclude that the selective potentiation of auditory corticostriatal synaptic strength is specific to the acquisition of the auditory task.

**Figure 3.**
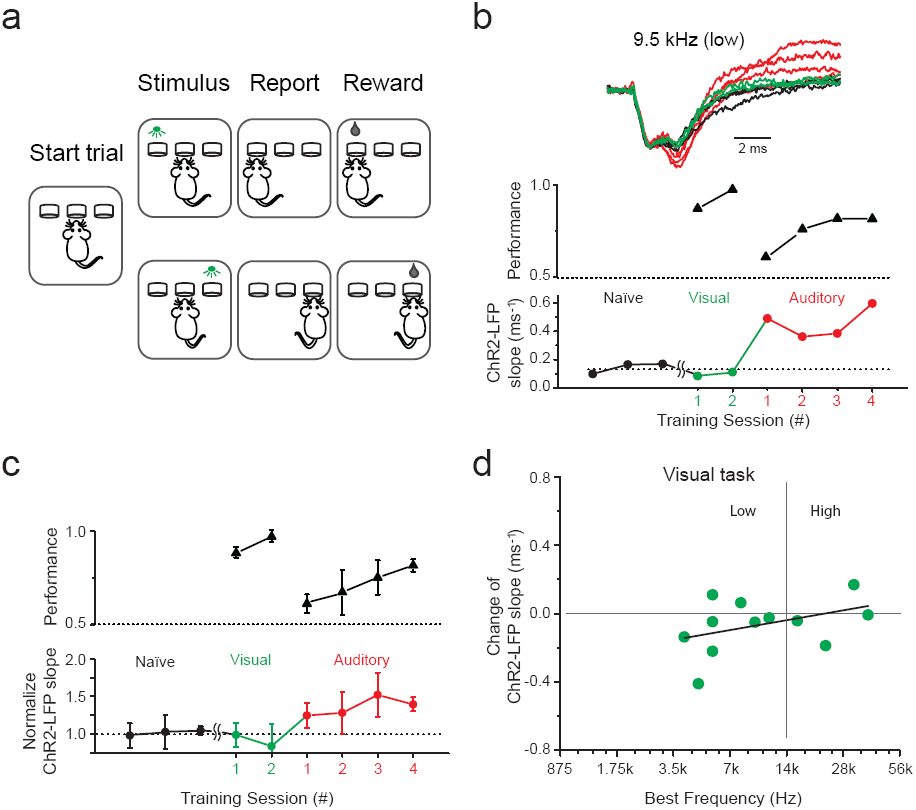
*Potentiation of ChR2-LFP slope is modality-specific.* (**a**) Visual 2-AFC task. **(b)** ChR2-LFP responses from an example auditory striatum site during visual and auditory task learning, analyzed as in Fig. 2a. Session 1 is defined as the first session in which the animal performed the full task (see Methods). (**c**) Population average of normalized ChR2-LFP slope during visual and auditory task training. (**d**) Visual training fails to potentiate auditory striatal input (12 recording sites from 4 rats; least squares regression, *p*=0.192).

The preferential potentiation *in vivo* of striatal sites tuned to low frequencies suggested that the pattern of potentiation within the striatum might be spatially organized. We therefore developed an *in vitro* brain slice preparation to examine this possibility. We first characterized the tonotopic organization of the auditory corticostriatal projection by injecting AAVs encoding either red or green fluorescent proteins at two different positions along the auditory cortical tonotopic axis. Cortical axons terminated in the striatum in distinct bands, with cortical projections tuned to high frequency sounds terminating more laterally in the auditory striatum and projections tuned to low frequency sounds more medially (Fig. 4a & S8). We next developed a protocol to assess the gradient of corticostriatal potentiation along the tonotopic axis, by recording ChR2-LFPs in coronal slices that preserve striatal tonotopy (Fig. 4b, see Methods). These recordings targeted left striatum, contralateral to the reward direction associated with low-frequency stimuli (LowRight; n=7 rats). For consistency across experiments, we used only a single slice from each animal, selected on the basis of striatal and hippocampal landmarks (see Methods). ChR2-LFPs in these slices showed a stereotyped waveform similar to that observed *in vivo,* and pharmacological dissection confirmed that the late component of the response was mediated mainly by AMPA-type glutamate receptors (Fig. 4c). In each slice we recorded the ChR2-LFP at between 8 and 16 sites (12.1 ± 2.1), and determined the gradient of ChR2-LFP along the tonotopic axis. Naïve rats showed no systematic difference in the strength of cortical input along the striatal tonotopic axis (Fig. S9). In contrast, in rats trained to associate low frequencies with rightward choices, the evoked corticostriatal response was strongest at medial (low frequency) sites and decreased laterally (Fig. 4d&e). This confirmed our observations *in vivo* and was consistent with the association to contralateral rewards. Thus the degree of corticostriatal synaptic potentiation induced by learning depended systematically on the position along the striatal tonotopic axis.

**Figure 4.**
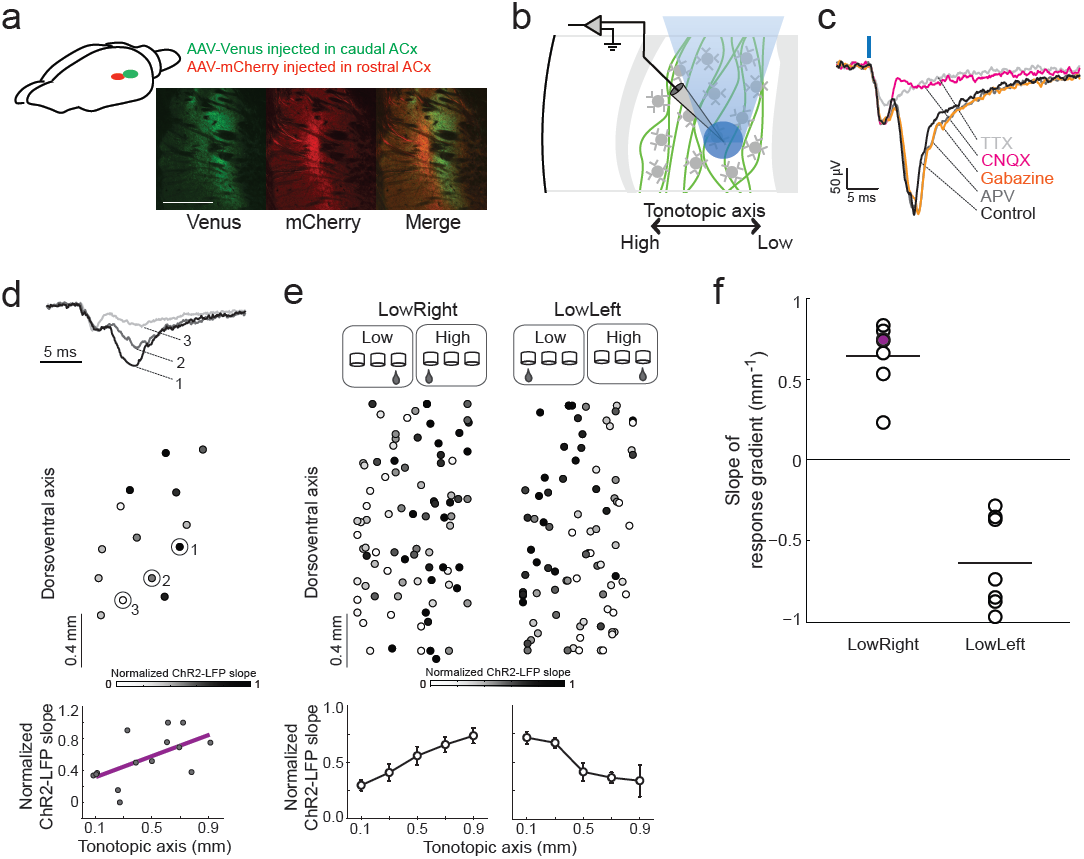
*Gradient of corticostriatal ChR2-LFP slopes encodes the association between stimulus and action.* (**a**) Tonotopy of projections from auditory cortex to striatum. Scale bar, 1 mm. **(b)** Recording paradigm. Light spot (blue) activates a subset of ChR2-expressing corticostriatal axons (green) near recording site. (**c**) Pharmacological dissection of corticostriatal ChR2-LFP responses in a striatal slice. (**d**) Normalized ChR2-LFP recorded at many sites within a striatal slice. Sample waveforms (1-3) shown above. ChR2-LFP slope increases with position along tonotopic axis (lower panel). (**e**) Population data for LowRight (n=7 rats) and LowLeft (n=7 rats). Error bars are s.e. **(f**) Gradient correctly identifies learned association in 14/14 individual rats (binomial test *p*=0.00006). Slope of example shown in **d** is indicated in purple. Black bars are the mean values for each group.

If the gradient of potentiation along the striatal tonotopic axis encodes the association between frequency and choice direction, then animals trained to make the opposite association should have a gradient of opposite sign. To test this we trained a new cohort of animals to associate low frequencies with leftward choices (LowLeft; n=7 rats). As predicted, the gradient in these animals was of similar magnitude but opposite in sign (Fig. 4e). There was no difference between these two training groups in ChR2-LFP across the orthogonal (dorsoventral) axis (*p*=0.22, paired *t*-test; Fig. S10). Thus the spatial gradient of corticostriatal potentiation induced by learning along the tonotopic axis depends on the training contingencies to which the animal is subjected.

Finally, we wondered whether the direction of the stimulus-response association could be inferred based on the sign of the ChR2-LFP gradient in individual animals. Remarkably, the training regimen (LowRight vs. LowLeft) could be correctly determined from the sign of gradient in single slice of every rat (14/14) tested (binomial test *p*=0.00006, Fig. 4f). Thus post-mortem study of corticostriatal efficacy can reliably reveal the training history of individual subjects.

Our results suggest a simple model of how the specific pattern of corticostriatal potentiation we observed might mediate task acquisition. In the LowRight task, training selectively potentiated corticostriatal synapses tuned to low frequencies between the left auditory cortex and the left auditory striatum (Fig. S11). Thus in behaving animals, low frequency tones would trigger stronger activation in the left auditory striatum and direct the animal to the right (contralateral) response port, possibly through the action of direct pathway medium spiny neurons^10^ that project ipsilaterally to the left substantia nigra pars reticulata and in turn to the superior colliculus^16^. On the other hand, in LowLeft trained animals, potentiation would cause the same stimulus to trigger stronger activation in right auditory striatum and direct the animal to the left response port. Although this model ignores much of the complexity of striatal circuitry, it offers a simple framework for understanding our results.

Previous work has demonstrated synaptic changes induced by learning^17-20^ or associated with reorganization of sensory maps^21-24^. Our results identify a locus of synaptic plasticity during the acquisition of a sensory discrimination task. We focused on auditory frequency discrimination, which allowed us to exploit the spatial organization of auditory corticostriatal connections to dissect the rules that govern corticostriatal plasticity during operant learning. Training selectively enhanced the strength of cortical inputs to establish an orderly gradient of corticostriatal synaptic strength across the striatal tonotopic axis.

The strengthening of a subset of connections, selected from a rich sensory representation, is reminiscent of several powerful models of learning^25-27^. In these models, even difficult nonlinear classification can be achieved by combining a high-dimensional representation of the stimulus—such as is found in the auditory cortex^28^ —through simple perceptron-like learning rules. We speculate that selective strengthening of appropriate corticostriatal synapses would allow animals to categorize a wide range of sensory stimuli—even those that are not mapped topographically in the striatum—and may reflect a general mechanism through which sensory representations guide the selection of motor responses.

## Materials and Methods

### Animals and viruses

Animal procedures were approved by the Cold Spring Harbor Laboratory Animal Care and Use Committee and carried out in accordance with National Institutes of Health standards. AAV-CAGGS-ChR2-Venus serotype 2/9 was packaged by the University of Pennsylvania Vector Core.

Long Evans male rats (Taconic Farm) were anaesthetized with a mixture of ketamine (50 mg kg^-1^) and medetomidine (0.2 mg kg^-1^), and injected with virus at 3-4 weeks old in the left auditory cortex. To cover most of the area and layers of the primary auditory cortex, 3-4 injections were made perpendicularly to brain surface at 1, 2, and 3 mm caudal to the temporoparietal suture, and 1 mm from the ventral edge. Each injection was made at three depths (500, 750 and 1000 μm), expelling approximately 200 nl of virus at each depth.

### Behavioural training

Rats were placed on a water deprivation schedule and trained to perform an auditory 2AFC task in a single-walled sound-attenuating training chamber as described previously^1^. Briefly, freely moving rats were trained to initiate a trial by poking into the center port of a three-port operant chamber, which triggered the presentation of a stimulus. Subjects then selected the left or right goal port. Correct responses were rewarded with water. The cloud-of-tones stimulus consisted of a stream of 30-ms overlapping pure tones presented at 100 Hz. The stream of tones continued until the rat withdrew from the center port. Eighteen possible tone frequencies were logarithmically spaced from 5 to 40 kHz. For each trial either the low stimulus (5 to 10 kHz) or high stimulus (20 to 40 kHz) was selected as the target stimulus, and the rats were trained to report low or high by choosing the correct side of port for water reward. In LowLeft task, the rats were required to go to the left goal port for water reward when the low stimulus was presented, and to the right goal port when high stimulus was presented. In LowRight task, the rats were required to go to the right goal port when the low stimulus was presented, and to the left goal port when the high stimulus was presented.

The pre-stimulus delay was drawn from exponential distribution with a mean of 300 ms. Early withdrawal from the center port before the onset of stimuli terminated the trial and a new trial was started. To complete a trial after exiting the center port, the animals were allowed up to 3 seconds to select a reward port. Typically, they made their choice within 300-700 ms. Error trials, where the rats reported to the wrong goal port after the presentation of the stimulus, were penalized with a 4 second time-out.

The intensity of individual tones was constant during each trial. To discourage rats from using loudness differences in discrimination, tone intensity was randomly selected on each trial from a uniform distribution between 45 and 75 dB (SPL) during training.

Implanted rats were water deprived and given free water for 1 hour every day before ChR2-LFP recording. These sessions were used to record baseline ChR2-LFP responses and defined as naïve sessions. Once a stable baseline was achieved, we began training subjects to perform the cloud-of-tones task. To introduce the subjects to the task structure, they were first trained (“direct mode”) to poke at the center port, which triggered the presentation of a stimulus and elicited water delivery from the corresponding goal port. Direct mode training was continued until a subject completed at least 150 trials in a single session (usually the first or second session). In subsequent sessions, defined as “session 1” in Figs. 2 and 3, the animal was trained on the “full task,” in which water was delivered only if the subject poked the correct goal port. For control subjects used in Fig. 3, subjects were trained in the direct mode with visual stimuli prior to introducing the full visual task, and then trained on the full auditory task.

### Tetrode recording and optogenetics

Custom tetrode and optic fiber arrays were assembled as described previously^1^. Each array carried 6 individually movable microdrives. Each microdrive consisted of one tetrode (4 polyimide-coated nichrome; wire diameter 12.7 μm; Kanthal Palm Coast) twisted together and gold-plated to an impedance of 0.3–0.5 MΩ at 1 kHz) and one optic fiber (62.5 μm diameter with a 50-μm core; Polymicro Technologies). The tetrode and fiber on the same microdrive were glued together, with the tips approximately 100 μm from each other.

To implant the tetrode/fiber array, rats were anaesthetized with a mixture of ketamine (50 mg kg^-1^) and medetomidine (0.2 mg kg^-1^) and placed in a stereotaxic apparatus. A craniotomy was made above the target area (2.5 to 3.5 mm from Bregma and 4 mm to 6 mm lateral from the midline). All rats were implanted in left hemisphere. The array was fixed in place with dental acrylic, and the tetrodes were lowered down to auditory striatum (3 to 5 mm from pia).

Electrical signals in auditory striatum were recorded using Neuralynx Cheetah 32-channel system and cheetah data acquisition software. For action potential recording, signals were filtered 600 to 6000 Hz. For local field potential recording, signals were filtered 10 to 9000 Hz. The rise time of the ChR2-LFP is relatively fast, so to preserve its dynamics we sought to stay as close to the raw data as possible. We re-analyzed the data in Fig 2a using two other choices of offline filter (median, 660 msec window & Butterworth lowpass 1-800 Hz). As shown in supplementary fig 12a&b, although filtering does affect the details of the ChR2-LFP shape (especially the earliest component), the results are qualitatively unchanged (Fig. S12c).

To determine the preferred sound frequency of recording sites, pure tones spanning from 1 kHz to 64 kHz were presented to rats before the start of behavioral training in a soundproof chamber for 100 ms every 2 s, in a random order at 30, 50 or 70 dB (SPL). The multi-unit baseline-subtracted firing rate in a window 5 to 55 ms after sound onset was compared with that in a window 0 to 50 ms before sound onset; only sites that significantly responded to sound were included. Firing rates in the window 5 to 55 ms after sound onset were computed for each frequency at 70 dB, and the peak of the resulting tuning curve was selected as the preferred frequency (Fig. S7).

For ChR2-LFP recording, 473 nm laser light was delivered through an FC/PC patch cord using a FiberPort Collimator (Thor Labs) to each implanted fiber individually. ChR2-LFP was recorded immediately after each training session. The laser power out of the patch cord was measured and adjusted to elicit an LFP with clear early and delayed components at each recording site (1-10 mW). For individual recording sites, laser power was adjusted slightly between days to maintain the early, presynaptic component of the LFP response at a consistent level (Fig. S13). Each light pulse was 100 μs in duration, presented at 1Hz, and each recording was an average of approximately 100 trials.

### *In vivo* pharmacology

To dissect the components of ChR2-LFP *in vivo*, rats were anaesthetized and placed in a stereotaxic apparatus. A single tetrode/fiber bundle was placed on a motorized manipulator (Sutter Instrument Company) and the tips of tetrode/fiber were guided to auditory striatum. Glass pipettes were used to deliver chemicals into auditory striatum. The pipettes filled with desired chemicals were carefully moved to penetrate through cortex and placed with the tips close to auditory striatum. Air pressure was slowly applied to inject the chemicals into tissue through a syringe that was connected to the pipette.

### Slice recording

Virus-injected and trained rats were anesthetized and decapitated, and the brains were transferred to a chilled cutting solution composed of (in mM) 110 choline chloride, 25 NaHCO_3_, 25 D-glucose, 11.6 sodium ascorbate, 7 MgCl_2_, 3.1 sodium pyruvate, 2.5 KCl, 1.25 NaH_2_PO_4_ and 0.5 CaCl_2_. Coronal slices (350 μm) were cut and transferred to artificial cerebrospinal fluid (ACSF) containing (in mM) 127 NaCl, 25 NaHCO_3_, 25 D-glucose, 2.5 KCl, 4 MgCl_2_, 1 CaCl_2_ and 1.25 NaH_2_PO_4_, aerated with 95% O_2_ 5% CO_2_.

To ensure maximal alignment across animals, only a single slice (350 μm thickness, between 2.5 mm and 2.9 mm from Bregma) per animal was used. Slices were incubated at 34 °C for 15–20 min and then kept at room temperature (22 °C) during the experiments. Local field potentials were recorded using Axopatch 200B amplifiers (Axons Instruments, Molecular Devices).

We delivered light pulses through a light guide microscope illumination system (Lumen Dynamics) modified to accept a blue laser (473 nm, Lasermate Group, CA, USA) in place of the lamp. The laser beam was focused onto the sample through the 60X objective during recordings, with illumination field of 350-μm diameter. Each light pulse was 500 μs at 1Hz, and each recording was an average of approximately 10 trials. Laser power was adjusted from site to site to maintain a similar level of axonal stimulation as judged by the amplitude of the early, presynaptic component of the ChR2-LFP response. To minimize the contribution of rundown on the estimation of the plasticity gradient within the striatal slice, recording locations were selected in randomly for each slice.

### Data analysis

All data were analyzed in MATLAB.

Behavior analysis only included completed trials. The percentage of correct trials for each animal in each session was computed using the last 200 trials of that session, unless the number of trials was less than 300, in which case only the last 100 trials were used.

Each measurement of *in vivo* ChR2-LFP was from a trace obtained by averaging across 70-100 trials (The slope measured from averaged trace is not different from the averaged slope from those of single traces, Fig. S14). Each average trace was normalized to the peak of the first component (around the time window between 0.5 ms to 1.2 ms after light stimulation, Fig. S2a), and the LFP slope was estimated by a linear regression fit of the rising phase of the second component (Fig. S2b, same time window was used for each recording site across sessions, the time window from site to site varied and was adjusted by eye for each site, ranged from 1.6 ms to 5 ms after light stimulation). The ChR2-LFP slope for each recording site across sessions was used as a measure of synaptic strength in subsequent analyses (Fig. 2a&b, Fig. 3b). The absolute change of the ChR2-LFP slope was used in Fig. 2d & 3d. In population analyses, normalized synaptic strength for each site was obtained by dividing the LFP slope values from all sessions by the mean of the ChR2-LFP slope values from naïve sessions (Fig. 2c & 3c).

For quantification of the ChR2-LFP in slice recording, the ChR2-LFP slope at each recording site was obtained in a manner similar to that *in vivo*: each averaged trace was normalized to the peak of the first component (around the time window between 1.5 ms to 4 ms after light stimulation), and the ChR2-LFP slope was estimated by a linear regression fit of the rising phase of the second component (around the time window between 5 ms to 8 ms after light stimulation, adjusted by eye for each slice). For each slice, the ChR2-LFP slopes across sites were rescaled from 0 to 1, with the smallest ChR2-LFP set to zero and the largest to 1. All recorded brain slices were aligned to a consensus brain slice. The positions of recording sites were measured from the aligned brain slices. Data from all the slices were pooled together for plotting the summary plot and for the quantitative analysis.

**Supplementary figure 1.**
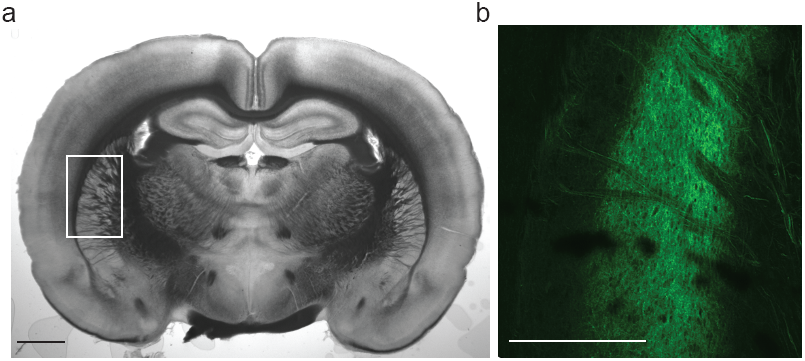
Corticostriatal projections from auditory cortex to striatum. **(a**) Coronal view for the location in the striatum that receives auditory cortical inputs. **(b**) Confocal image of auditory cortical axon terminals expressing ChR2-Venus in the striatum. Scale bars: 2 mm.

**Supplementary figure 2.**
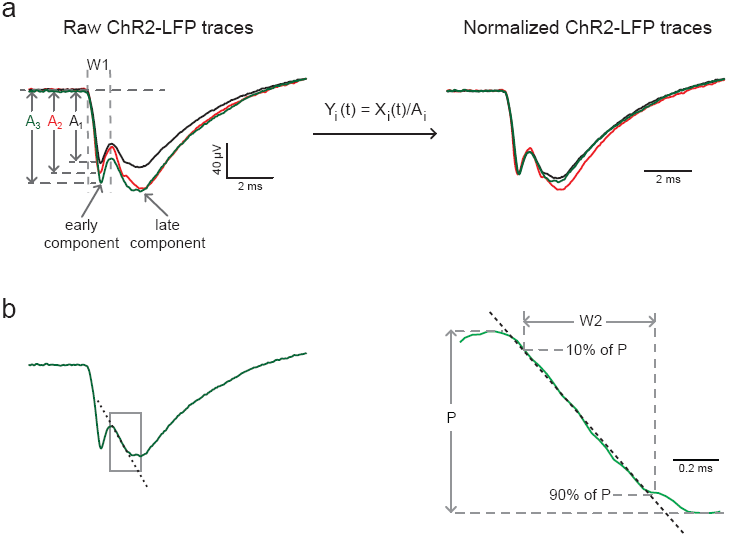
Normalization method and slope measurement for ChR2-LFP. **(a**) Raw ChR2-LFP traces (left panel) were normalized to the amplitude of their corresponding early component (A_i_). The normalization factor A_i_ was determined as the peak of the raw trace in the time window (W1) between 0.5 ms to 1.2 ms after light stimulation onset. **(b**) The rising phase of the late component of ChR2-LFP (in a time window W2 defined by rise from 10% to 90% of the peak P) was fit linearly, and the slope of the fit was used for the quantification of ChR2-LFP.

**Supplementary figure 3.**
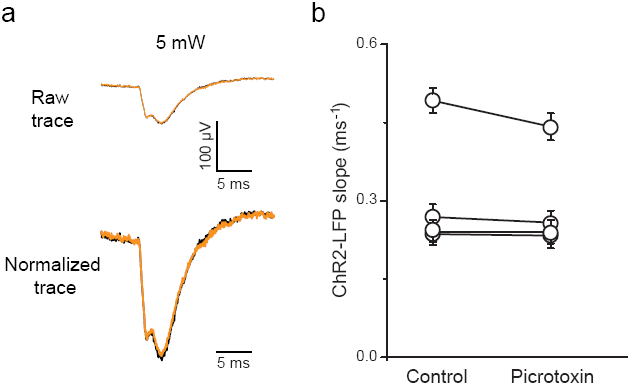
GABAergic block by picrotoxin had no effect ChR2-LFP slope *in vivo*. **(a**) ChR2-LFP before (black traces) and after (orange traces) picrotoxin application (20 mM, 5 ul). Raw traces are averaged traces from 60-80 trials at each condition (upper row). Normalized traces are raw traces normalized to their peaks of first components (as illustrated in fig S2). **(b**) Slopes measured from normalized traces in control and picrotoxin conditions for each recording before and after picrotoxin application (*p*=0.8, paired signed-rank test). Data in **b** are presented as mean ± s.e..

**Supplementary figure 4.**
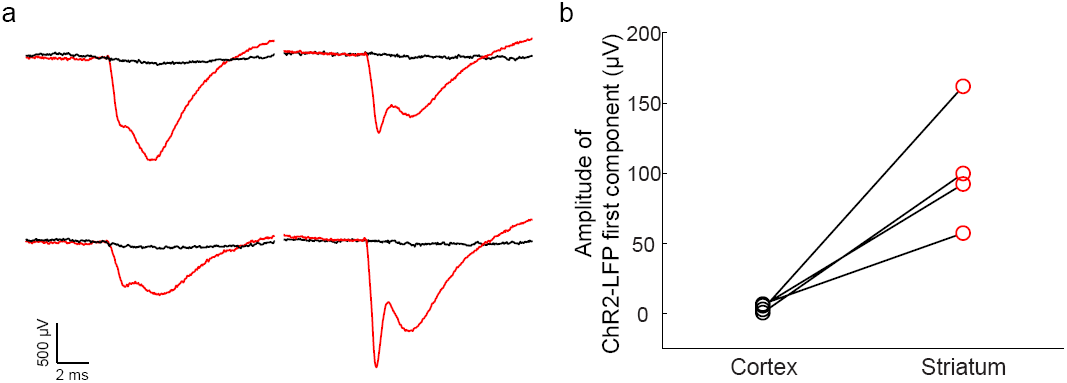
ChR2-LFP depends on presence of ChR2-expressing axons. **(a**) Four independent recordings in the auditory striatum (red traces) which receives auditory cortical input (ChR2-expressing axon terminals are present), and the overlying somatosensory cortex (black traces) which lacks auditory cortical input (ChR2-expressing axon terminals are absent). Each pair of recordings is from the same tetrode/fiber bundle. The recordings indicate that the light artifact is negligible under our conditions. **(b**) Comparison of the first component amplitude from each recording pair.

**Supplementary figure 5.**
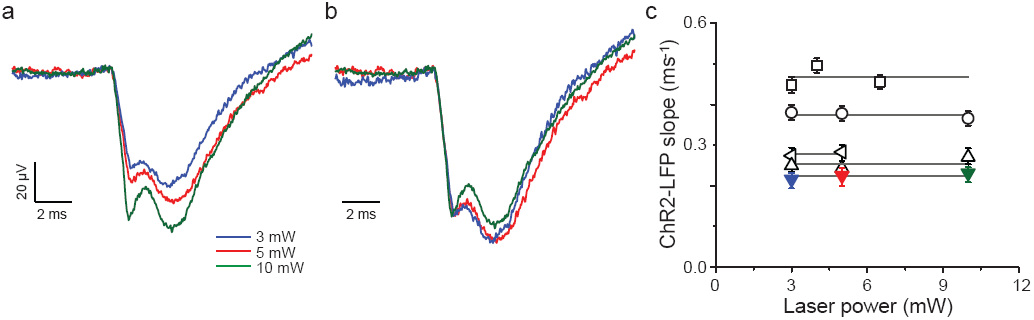
Normalization procedure corrects for changes in light power *in vivo*. **(a**) One example of ChR2-LFP recorded at different light levels. **(b**) Normalized ChR2-LFP the same example in a. **(c**) Slopes from 5 example recordings across 1-10 mW light level range (colored symbols are examples shown in **a** & **b**). Grey lines are drawn from the mean values of each group.

**Supplementary figure 6.**
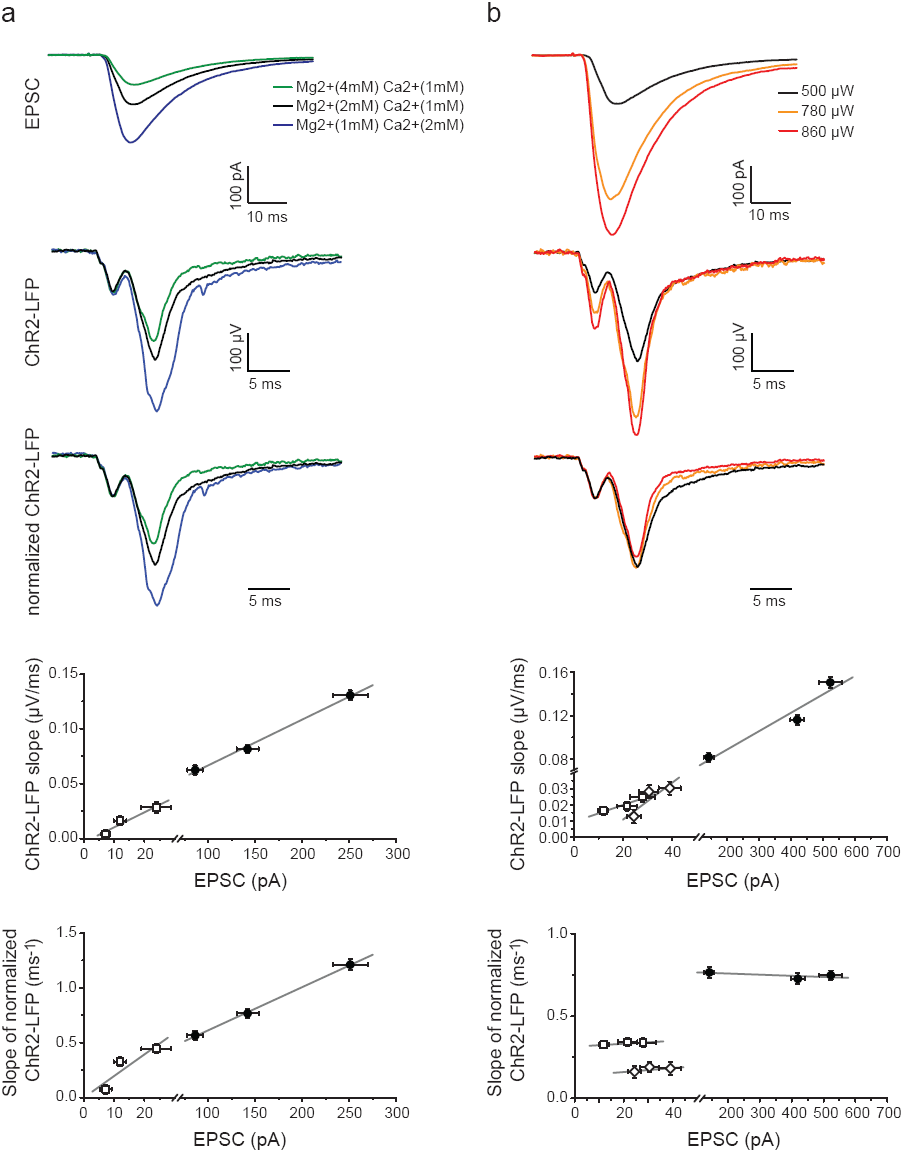
The ChR2-LFP recorded extracellularly mirrors the ChR2-EPSC recorded from single neurons. The synaptic response was manipulated by either changing synaptic release probability or the number of fibers recruited with a light pulse. **(a**) The concentration of divalent cations in the extracellular solution was varied to change synaptic release probability. Paired *in vitro* whole-cell voltage clamp recording example (EPSC, first row) and LFP (second and third rows) at different external divalent ion concentrations. Slopes of LFP measured from raw traces (fourth row, solid circles) and normalized traces (fifth row, solid circles) changed linearly with EPSC amplitudes (R^2^ = 0.96 & 0.99 respectively). Grey lines in fourth and fifth rows are linear regression fit for each recording pair (R^2^ = 0.94 & 0.81 for squares in fouth and fifth row respectively). **(b**) We varied the light level to change the number of ChR2-expressing axons recruited by a light flash. Paired in vitro whole-cell voltage clamp recording (EPSC, first row) and LFP (second and third rows) at different light levels. Slopes of the LFPs measured from raw traces (fourth row, solid circles) changed monotonically with EPSC amplitudes (R^2^ = 0.94 for solid circles, R^2^ = 0.91 for squares and 0.77 for diamonds). Slopes of the normalized LFPs remain constant at different light levels (fifth row, solid circles for the example shown in first and second rows). The normalization procedure thus minimizes fluctuations in the response arising from artifactual changes in the number of recruited fibers, but preserves changes arising from actual increases or decreases in synaptic efficacy.

**Supplementary figure 7.**
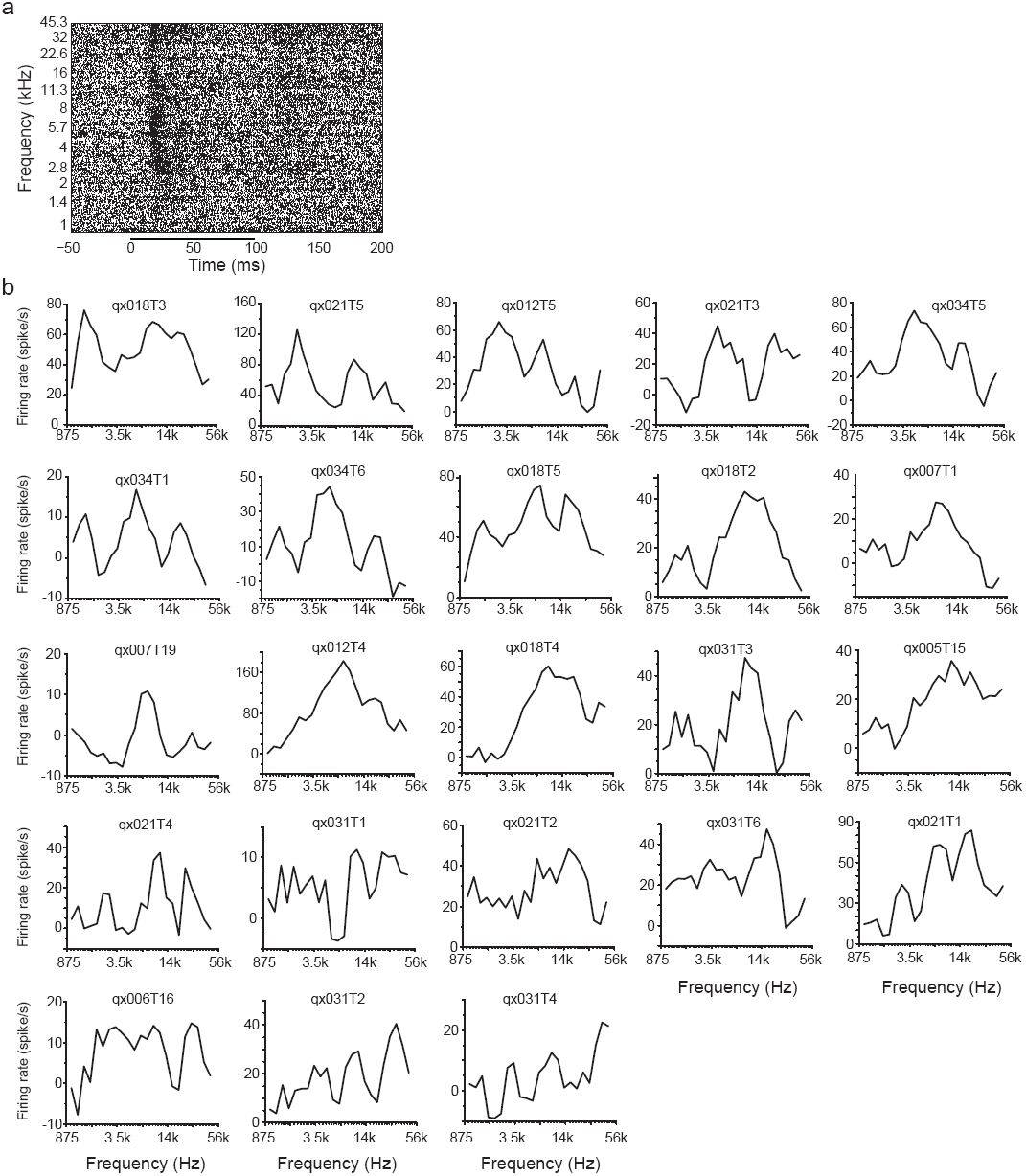
Multiunit responses in the auditory striatum are tuned to frequency. **(a**) Raster plot of multi-unit recordings from one example site in the auditory striatum. Pure tones with different frequencies were presented between 0 and100 ms. Each frequency was presented in a random order for 20-30 trials. **(b**) Tuning curves for individual recording sites that are included in Figure 2d.

**Supplementary figure 8.**
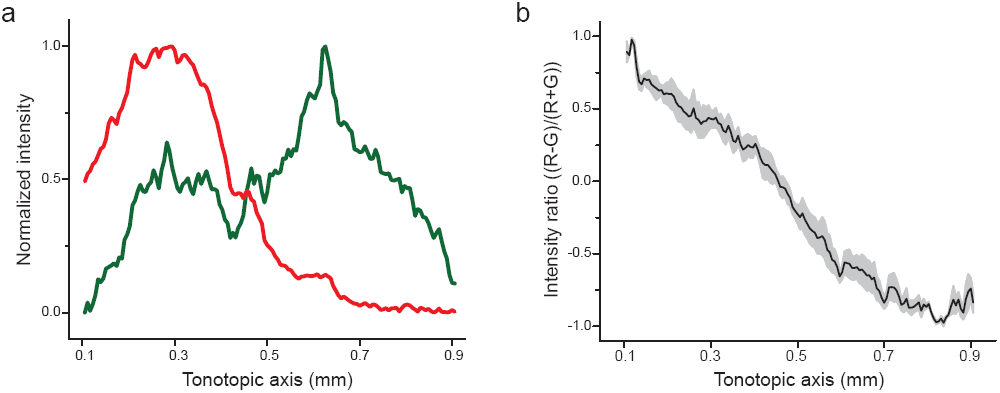
Quantification of corticostriatal projection. **(a**) Normalized red and green fluorescence intensities measured across the tonotopic axis from image shown in Figure 4a. **(b**) Average red/green intensity ratio across the tonotopic axis, n = 3 sections from 2 rats.

**Supplementary figure 9.**
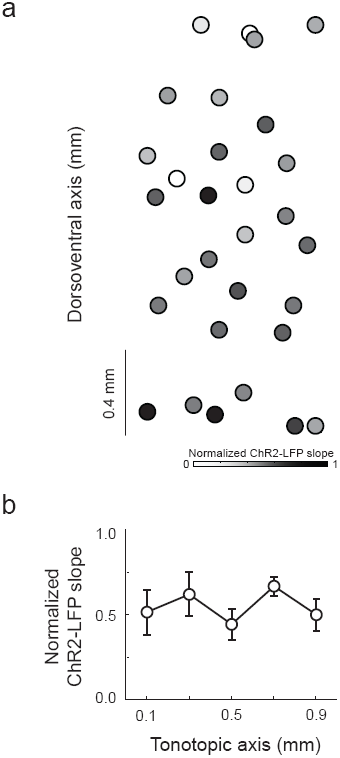
No corticostriatal ChR2-LFP slope gradient across tonotopic axis in naïve rats. **(a)** ChR2-LFP slope map from 3 striatal slices (n = 3 rats). **(b**) Quantification of ChR2-LFP slope across tonotopic axis. Data are presented as mean ± s.e..

**Supplementary figure 10.**
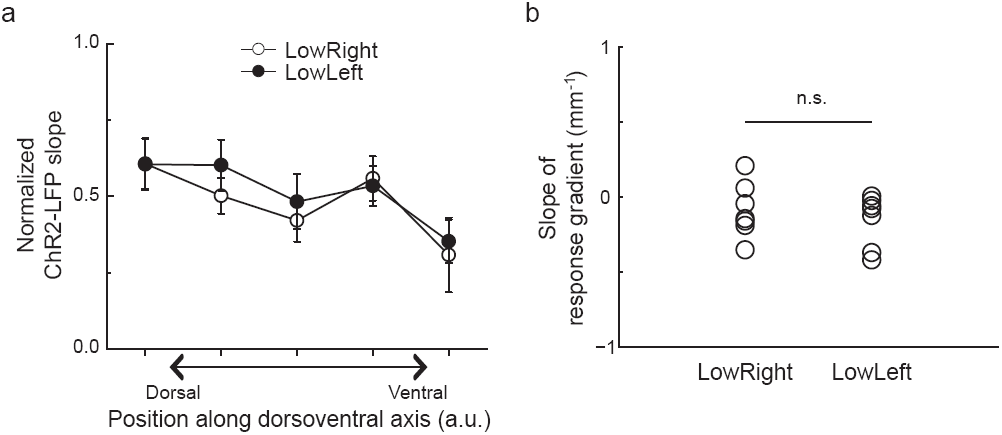
Gradient of ChR2-LFP across the dorsoventral (non-tonotopic) axis showed no difference between two training groups. **(a)** Averaged ChR2-LFP slopes with position along tonotopic axis for LowRight and LowLeft (7 rats from each group). **(b)** Individual gradients of ChR2-LFP across dorsoventral aixs from LowRight and LowLeft groups (*p*=0.22, paired *t*-test).

**Supplementary figure 11.**
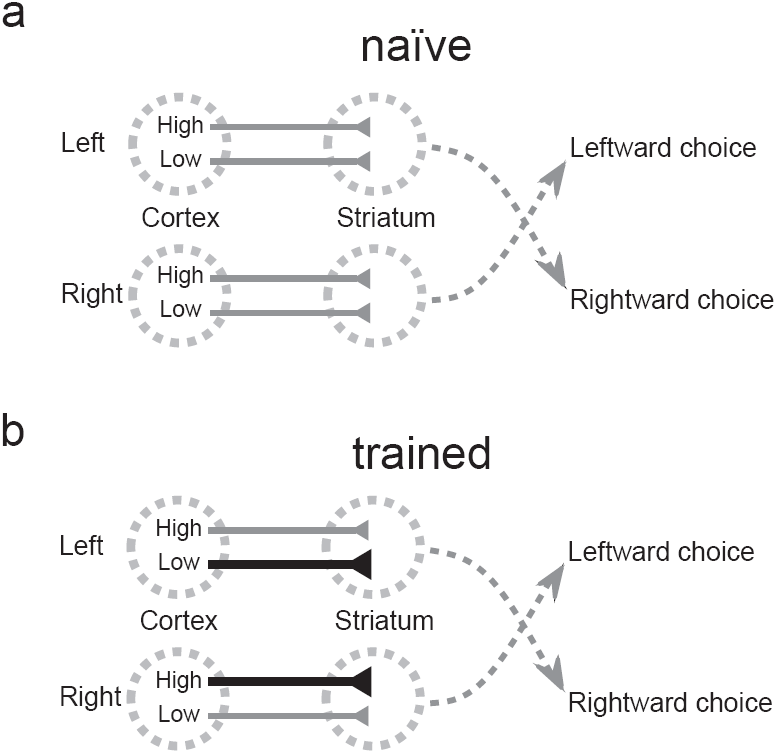
Model showing how corticostriatal potentiation could mediate task acquisition. **(a)** In the naïve rat, the strength of corticostriatal connections does not depend on their frequency preference. **(b)** Training to associate low stimuli with rightward choices and high stimuli with leftward choices (LowRight) selectively potentiates corticostriatal synapses tuned to low frequencies in the left hemisphere and corticostriatal synapses tuned to high frequencies in the right hemisphere. Thus in the trained rat, low stimuli drive rightward choices and high stimuli drive leftward choices.

**Supplementary figure 12.**
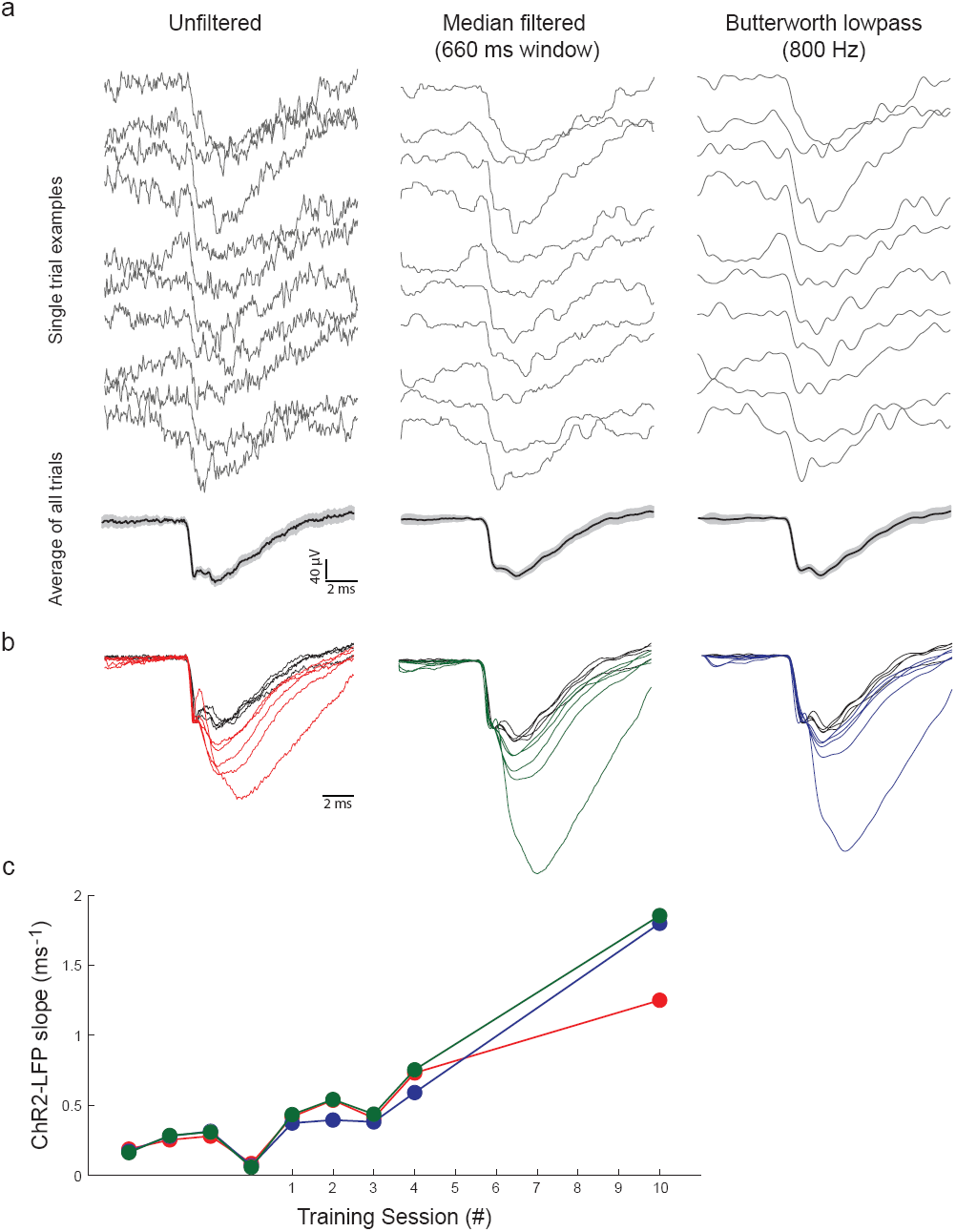
ChR2-LFP measurement is not affected by different filter settings. **(a)** Single trial (upper rows) and average (bottom row) examples of unfiltered, median filtered and Butterworth lowpass filtered responses. Average traces are presented as mean values (black traces) with 95% confidence (grey shading). **(b)** ChR2-LFP examples in **Figure 2a** with different filter settings. **(c**) ChR2-LFP measurements from examples shown in **Figure 2a** at different filter settings.

**Supplementary figure 13.**
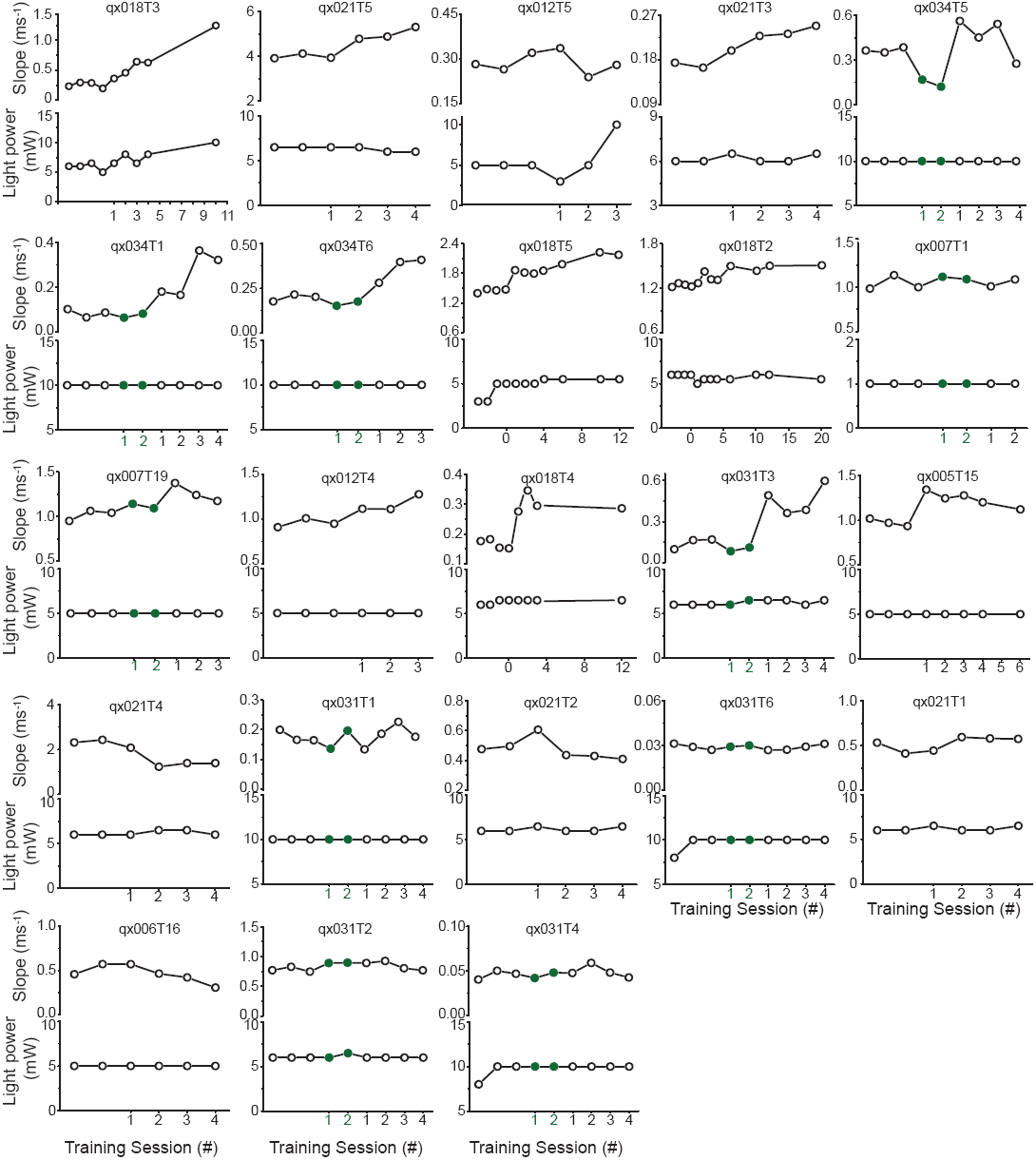
Changes of ChR2-LFP across training sessions are not related to light level adjustments. Each panel consists two plots for the same recording site: slopes across sessions (upper plot) and light levels across sessions (lower plots). Green colored sessions represent visual task training, and black colored sessions represent auditory task training.

**Supplementary figure 14.**
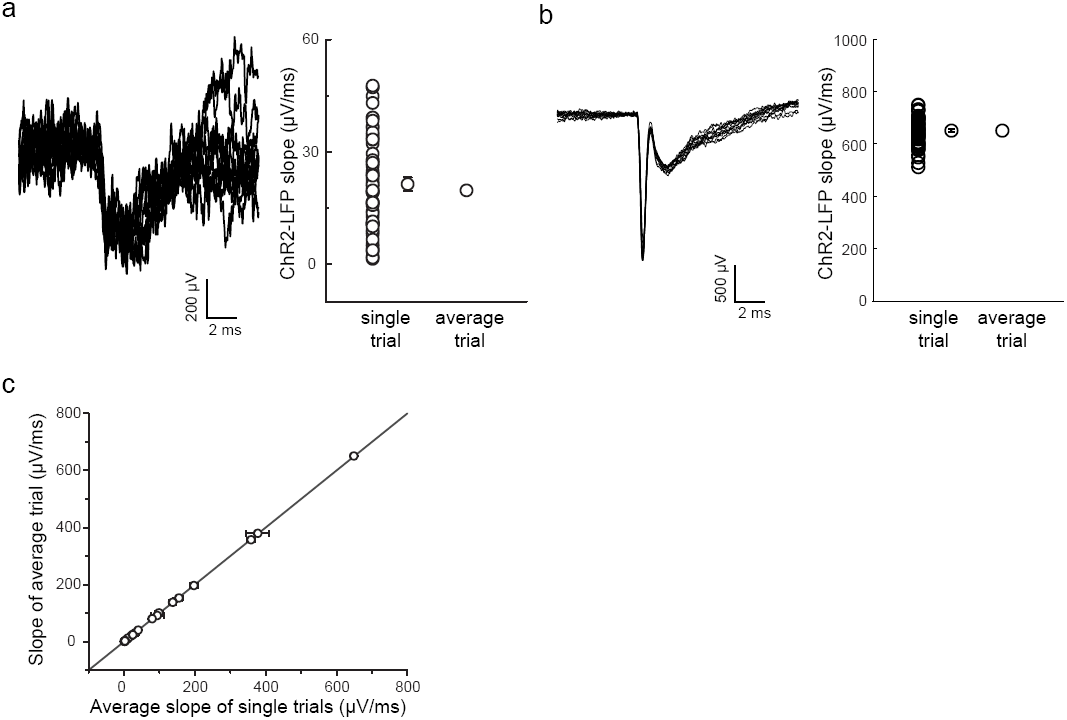
Slopes measured from single trials and averaged traces are similar. **(a)** One noisy example for single trials traces (left panel) and slopes measured from individual traces and averaged trace (right panel). **(b)** One less-noisy example. **(c**) Comparison of averaged slopes from single trial measurement and slopes measured from averaged traces.

## Acknowledgments

We thank Barry Burbach for invaluable technical help. AAV-CAGGS-ChR2-Venus was provided by K. Svoboda. This work was supported by grants from the National Institutes of Health (A.M.Z.).

## References

1. Znamenskiy, P. & Zador, A. M. Corticostriatal neurons in auditory cortex drive decisions during auditory discrimination. Nature 497, 482–485, DOI:10.1038/nature12077 (2013).

2. Salzman, C. D., Britten, K. H. & Newsome, W. T. Cortical microstimulation influences perceptual judgements of motion direction. Nature 346, 174–177, DOI:10.1038/346174a0 (1990).

3. Roitman, J. D. & Shadlen, M. N. Response of neurons in the lateral intraparietal area during a combined visual discrimination reaction time task. The Journal of neuroscience : the official journal of the Society for Neuroscience 22, 9475–9489 (2002).

4. Uchida, N. & Mainen, Z. F. Speed and accuracy of olfactory discrimination in the rat. Nat Neurosci 6, 1224–1229, DOI:10.1038/nn1142 (2003).

5. Felsen, G. & Mainen, Z. F. Neural substrates of sensory-guided locomotor decisions in the rat superior colliculus. Neuron 60, 137–148, DOI:10.1016/j.neuron.2008.09.019 (2008).

6. Erlich, J. C., Bialek, M. & Brody, C. D. A cortical substrate for memory-guided orienting in the rat. Neuron 72, 330–343, DOI:10.1016/j.neuron.2011.07.010 (2011).

7. Raposo, D., Sheppard, J. P., Schrater, P. R. & Churchland, A. K. Multisensory decision-making in rats and humans. J Neurosci 32, 3726–3735, DOI:10.1523/JNEUROSCI.4998-11.2012 (2012).

8. Brunton, B. W., Botvinick, M. M. & Brody, C. D. Rats and humans can optimally accumulate evidence for decision-making. Science 340, 95–98, DOI:10.1126/science.1233912 (2013).

9. Thompson, J. A. & Felsen, G. Activity in mouse pedunculopontine tegmental nucleus reflects action and outcome in a decision-making task. J Neurophysiol 110, 2817–2829, DOI:10.1152/jn.00464.2013 (2013).

10. Tai, L., Lee, A., Benavidez, N., Bonci, A. & Wilbrecht, L. Transient stimulation of distinct subpopulations of striatal neurons mimics changes in the value of competing actions. Nature Neuroscience 15, in press (2012).

11. Barnes, T. D., Kubota, Y., Hu, D., Jin, D. Z. & Graybiel, A. M. Activity of striatal neurons reflects dynamic encoding and recoding of procedural memories. Nature 437, 1158–1161, DOI:10.1038/nature04053 (2005).

12. Schultz, W. & Dickinson, A. Neuronal coding of prediction errors. Annu Rev Neurosci 23, 473–500, DOI:10.1146/annurev.neuro.23.1.473 (2000).

13. Reynolds, J. N., Hyland, B. I. & Wickens, J. R. A cellular mechanism of reward-related learning. Nature 413, 67–70, DOI:10.1038/35092560 35092560 [pii] (2001).

14. Yin, H. H. et al. Dynamic reorganization of striatal circuits during the acquisition and consolidation of a skill. Nature neuroscience 12, 333–341, DOI:10.1038/nn.2261 (2009).

15. Malenka, R. C. & Kocsis, J. D. Presynaptic actions of carbachol and adenosine on corticostriatal synaptic transmission studied in vitro. J Neurosci 8, 3750–3756 (1988).

16. Hikosaka, O., Takikawa, Y. & Kawagoe, R. Role of the basal ganglia in the control of purposive saccadic eye movements. Physiol Rev 80, 953–978 (2000).

17. Carew, T. J., Walters, E. T. & Kandel, E. R. Associative learning in Aplysia: cellular correlates supporting a conditioned fear hypothesis. Science 211, 501–504 (1981).

18. Rioult-Pedotti, M. S., Friedman, D. & Donoghue, J. P. Learning-induced LTP in neocortex. Science 290, 533–536 (2000).

19. Rumpel, S., LeDoux, J., Zador, A. & Malinow, R. Postsynaptic receptor trafficking underlying a form of associative learning. Science 308, 83–88 (2005).

20. Whitlock, J. R., Heynen, A. J., Shuler, M. G. & Bear, M. F. Learning induces long-term potentiation in the hippocampus. Science 313, 1093–1097, DOI:10.1126/science.1128134 (2006).

21. Finnerty, G. T., Roberts, L. S. & Connors, B. W. Sensory experience modifies the short-term dynamics of neocortical synapses. Nature 400, 367–371. (1999).

22. Trachtenberg, J. T. et al. Long-term in vivo imaging of experience-dependent synaptic plasticity in adult cortex. Nature 420, 788–794, DOI:10.1038/nature01273 (2002).

23. Froemke, R. C., Merzenich, M. M. & Schreiner, C. E. A synaptic memory trace for cortical receptive field plasticity. Nature 450, 425–429 (2007).

24. Hofer, S. B., Mrsic-Flogel, T. D., Bonhoeffer, T. & Hubener, M. Experience leaves a lasting structural trace in cortical circuits. Nature 457, 313–317, DOI:10.1038/nature07487 (2009).

25. Maass, W., Natschlager, T. & Markram, H. Real-time computing without stable states: a new framework for neural computation based on perturbations. Neural computation 14, 2531–2560, DOI:10.1162/089976602760407955 (2002).

26. Sussillo, D. & Abbott, L. F. Generating coherent patterns of activity from chaotic neural networks. Neuron 63, 544–557, DOI:10.1016/j.neuron.2009.07.018 (2009).

27. Cortes, C. & Vapnik, V. Support-vector networks. Mach Learn 20, 273–297, DOI:10.1007/BF00994018 (1995).

28. Hromadka, T., Deweese, M. R. & Zador, A. M. Sparse representation of sounds in the unanesthetized auditory cortex. PLoS biology 6, e16, DOI:10.1371/journal.pbio.0060016 (2008).

